# Sexual dimorphism and the effect of wild introgressions on recombination in cassava (*Manihot esculenta* Crantz) breeding germplasm

**DOI:** 10.1101/2021.09.22.461431

**Authors:** Ariel W. Chan, Seren S. Villwock, Amy L. Williams, Jean-Luc Jannink

**Affiliations:** Section of Plant Breeding and Genetics, School of Integrative Plant Sciences, Cornell University, Ithaca, NY 14853; Department of Biological Statistics and Computational Biology, Cornell University, Ithaca, NY 14853; RW Holley Center for Agriculture and Health, United States Department of Agriculture -- Agricultural Research Service, School of Integrative Plant Sciences, 258 Emerson Hall, Cornell University, Ithaca, NY 14853

## Abstract

Recombination has essential functions in meiosis, evolution, and breeding. The frequency and distribution of crossovers dictate the generation of new allele combinations and can vary across species and between sexes. Here, we examine recombination landscapes across the 18 chromosomes of cassava (*Manihot esculenta* Crantz) with respect to male and female meioses and known introgressions from the wild relative *Manihot glaziovii*. We used SHAPEIT2 and duoHMM to infer crossovers from genotyping-by-sequencing (GBS) data and a validated multi-generational pedigree from the International Institute of Tropical Agriculture (IITA) cassava breeding germplasm consisting of 7,020 informative meioses. We then constructed new genetic maps and compared them to an existing map previously constructed by the International Cassava Genetic Map Consortium (ICGMC). We observed higher recombination rates in females compared to males, and lower recombination rates in *M. glaziovii* introgression segments on chromosomes 1 and 4, with suppressed recombination along the entire length of the chromosome in the case of the chromosome 4 introgression. Finally, we discuss hypothesized mechanisms underlying our observations of heterochiasmy and crossover suppression and discuss the broader implications for plant breeding.

## INTRODUCTION

Meiotic recombination plays essential roles in evolution and breeding by creating new combinations of existing alleles, which generates genomic diversity that can be selected upon in a population (Barton and Charlesworth 1998). In the context of meiosis, crossing over aids in homology recognition and ensures proper segregation of homologous chromosomes to prevent aneuploidy (Moore and Orr-Weaver 1997). Recombination also serves as an important breeding tool, as its rate dictates the resolving power of quantitative trait mapping, the precision of allele introgression, and ultimately the ability to combine favorable alleles in the same haplotype for generating improved varieties (Mercier et al. 2015).

While recombination rates can vary among and within taxa, the variation appears to be tightly constrained by both an upper and lower bound (Ritz et al. 2017). In most species, there is one obligatory crossover per tetrad to prevent aneuploidy, which explains the lower bound (Wang et al. 2015). The reasons for an upper bound on crossover number, however, are less obvious. One plausible explanation is that limiting crossovers confers an evolutionary advantage by preserving favorable combinations of alleles residing on the same haplotype (Ritz et al. 2017).

The distribution of crossovers along chromosomes is not random and is influenced by chromosome features such as chromatin structure, gene density, and nucleotide composition (Dluzewska et al. 2018). The occurrence of a crossover at one location also reduces the likelihood that another crossover will occur in close proximity (Sturtevant 1915; Mercier et al. 2015). This nonrandom placement of crossovers, known as crossover interference, results in a pattern where recombination events appear more evenly spaced than would be expected by random chance (Foss et al. 1993). Interference may serve as a biological mechanism to ensure that every pair of homologous chromosomes undergoes at least one crossover event, which is necessary for proper disjunction (Otto and Payseur 2019).

In many species, crossover frequency and distribution along chromosomes differs between female and male meiosis, a phenomenon referred to as heterochiasmy (Lenormand and Dutheil 2005). The direction and degree of these differences are typically species-specific. The most extreme are cases in which one of the two sexes lacks meiotic recombination entirely; for example, male *Drosophila melanogaster* do not recombine during meiosis (Morgan 1910). In plants, the ratio of male to female recombination has been found to vary from 0.6 to 1.3 (Lenormand and Dutheil 2005). In wild-type *Arabidopsis thaliana,* male recombination is higher than female recombination, while the opposite was recently found in mutant lines with increased recombination (Fernandes et al. 2018). In a plant breeding context, heterochiasmy leads to an altered probability of generating a favorable recombination depending on the direction of a cross.

To date, recombination landscapes have not yet been well-characterized in cassava *(Manihot esculenta).* Cassava is a root crop cultivated in the tropics, a staple carbohydrate-rich food for hundreds of millions of people and a particularly important food security resource for small-holder farmers (http://faostat.fao.org). Recent genomic selection efforts have generated a large amount of genomic data, which also makes cassava a useful model for other tubers and clonally propagated crops (Ceballos et al. 2012; Wolfe et al. 2017). Cassava is a diploid organism with an estimated genome size of approximately 772 Mb spread across 18 chromosomes (Awoleye et al. 1994), with the reference genome spanning 582.28 Mb (Bredeson et al. 2016). The International Cassava Genetic Map Consortium (ICGMC) generated a consensus genetic map of cassava that combines 10 mapping populations, consisted of one self-pollinated cross and nine biparental crosses (14 parents total; 3,480 meioses) (ICGMC 2015). The genetic map is 2,412 cM in length and organizes 22,403 genotyping-by-sequencing (GBS) markers.

An important feature of the cassava genome in some populations is the presence of two large introgressions from the wild relative *Manihot glaziovii*. In the 1930’s, breeders crossed cassava with *M. glaziovii* to incorporate cassava mosaic disease resistance, and these hybrids were key founders of breeding germplasm (Hahn et al. 1980; Wolfe et al. 2019). Wolfe *et al.* 2019 detected large *M. glaziovii* introgressions prevalent in African cassava populations on chromosome 1, spanning from 25 Mb to the end of the chromosome, and on chromosome 4 from 5 Mb to 25 Mb. *M. glaziovii* and *M. esculenta* diverged approximately 2-3 million years ago, and have 2.2% homozygous differences at genotyped positions (Bredeson et al. 2016). The *M. glaziovii* introgressions are thought to contribute both beneficial alleles and deleterious load, are associated with strong linkage disequilibrium, and have been increasing in frequency although maintained in the heterozygous state in the IITA genomic selection program (Wolfe et al. 2019). Therefore, further investigation into the effect of *M. glaziovii* introgressions on recombination is needed to understand the implications of the introgression for cassava breeding.

Here, we demonstrate the application of SHAPEIT2 and duoHMM (O’Connell et al. 2014) to detect crossover events and characterize recombination landscapes across the cassava genome. Using a multi-generational cassava breeding pedigree and associated GBS data from the International Institute of Tropical Agriculture (IITA), we identified and validated informative parent-offspring duos and trios, phased parental haplotypes, and inferred recombination events between SNP intervals. In this context, we use both the terms “recombination” and “crossover” to refer to meiotic crossovers inferred from patterns of switched parental haplotypes observed in the progeny, though we note that not all crossovers result in a detectable exchange of polymorphisms, and that recombination can also occur due to homologous repair or gene conversion. We used the inferred crossovers to construct new genetic maps and compared them to the existing ICGMC composite map. We then examined crossover frequency and distribution across the genome with respect to sex and *M. glaziovii* introgression status. Finally, we discuss the implications of our observations for plant breeding.

## MATERIALS AND METHODS

### The IITA germplasm population structure

This study analyzed germplasm from the genomic selection program at IITA from 2012-2015 as part of the Next Generation Cassava Breeding Project (“NextGen”; nextgencassava.org). The IITA pedigree consists of 7,432 unique individuals from four breeding populations, originating from the Genetic Gain collection previously described in Okechukwu and Dixon 2008: Genetic Gain (GG; *n* = 494), TMS13 (*n* = 2,334), TMS14 (*n* = 2,515), or TMS15 (*n* = 2,089). Of the 494 GG individuals, 236 individuals are founders and the remaining 258 are the progeny of within-population GGxGG crosses. TMS13, TMS14, and TMS15 successively originated from crosses among and between the previous populations as illustrated in Fig. 1. The selection of parents used to generate each population are described in Wolfe *et al.* 2017.

**Figure 1.**
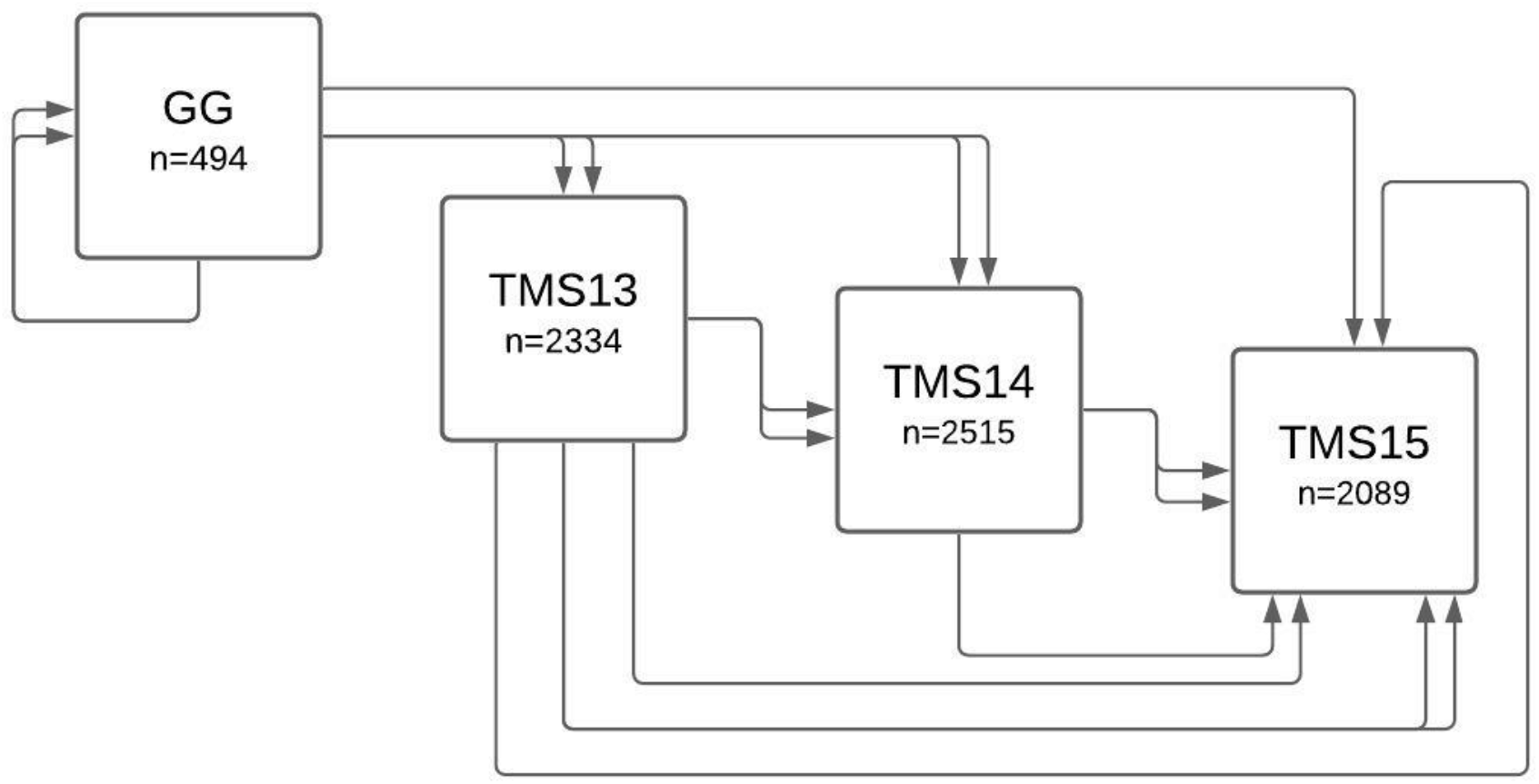
Diagram of the IITA pedigree structure. Population size and ancestry of the four breeding populations in the IITA pedigree. Arrows represent parentage relationships, where a pair of adjacent arrows represents two parents used in a cross.

### GBS genotyping and validation

The breeding populations were genotyped with GBS as described in Wolfe *et al.* 2017. GBS data consisting of 22,403 markers with an average depth of 7x was available through NextGen for 7,294 of the 7,432 individuals (*n_GG_* = 366, *n_TMS13_* = 2330, *n_TMS14_* = 2509, and *n_TMS15_* = 2089). Filters were applied to remove sites with more than 70% missing data, individuals with more than 80% missing data, and sites with a mean depth across all samples greater than 120 to avoid spurious genotype calls within repeat regions.

Some accessions in the population had more than one GBS record due to multiple sequencing events. Before merging the data, the R package BIGRED was used to verify the identity of putative technical replicates. BIGRED uses a Bayesian model to infer which of the samples originated from an identical genotypic source, as described in Chan *et al.* 2018. Putative replicates with unambiguous BIGRED results were inferred to be true replicates and were merged. Those for which BIGRED returned a source vector with no clear majority were ambiguous, so the samples were excluded from future analyses. Table 1 summarizes the number of individuals in each group with more than one GBS record that could be validated as replicates with BIGRED.

**Table 1.** Summary of GBS data records for each breeding group.

| Group ID | Individuals with available GBS data | Individuals with multiple GBS records | Replicated individuals resolved with BIGRED | Final number of individuals included for analysis |
| --- | --- | --- | --- | --- |
| GG | 366 | 189 | 168 | 345 |
| TMS13 | 2330 | 156 | 146 | 2320 |
| TMS14 | 2509 | 62 | 59 | 2506 |
| TMS15 | 2089 | 0 |  | 2089 |

The number of individuals with available GBS data in each breeding group. Of those with multiple GBS records, replicates that could not be unambiguously verified as identical with BIGRED were excluded from analysis.

### Validation of pedigree records using AlphaAssign

To validate pedigree information, the parentage assignment algorithm AlphaAssign was used to infer parents from GBS data. As described in Whalen *et al.* 2019, AlphaAssign uses the genotypes of a target individual and a known parent (if available) to calculate the posterior probability distribution of expected genotypes for its relatives and classify candidate individuals as one of four possible relationships: parent, full-sibling of a parent, half-sibling of a parent, or unrelated to the target individual. The list of candidate parents for each breeding group was based on the generation of that population as described above. For example, for TMS14 target individuals, GG and TMS13 individuals were listed as candidate parents. Founders were excluded as target individuals. To simplify the computations and to filter for sites in linkage disequilibrium, allelic depth data from 1,000 sites were sampled randomly across the 18 chromosomes such that no two sites fell within 20 kb from one another. The choice of this site count was based on the simulations of Whalen *et al.* 2019 which found this number of markers to be sufficient for accurate parentage assignment at 5x coverage.

Since AlphaAssign evaluates pairwise relationships, the validation procedure was carried out twice to infer both parents of individuals in the pedigree. In the first run, no prior pedigree information was given to the algorithm such that all calculations involved the use of a ‘dummy parent’ with genotype probabilities calculated using estimated allele frequencies and assuming Hardy-Weinberg Equilibrium (Whalen *et al.* 2019). For each target individual, the candidate individual with the highest score statistic was listed as the parent in an inferred pedigree. Inferred parents were then input as prior known parents for a second run to identify the other parent. The AlphaAssign-inferred pedigree was then compared with IITA’s existing pedigree to validate the listed parents. The 5,479 individuals with one or both listed parents successfully validated by AlphaAssign were considered useable as duos or trios for further analysis. Table 2 summarizes the number of individuals with validated parents from each breeding group.

**Table 2.** Summary of pedigree validation using AlphaAssign.

|  | GG | TMS13 | TMS14 | TMS15 |
| --- | --- | --- | --- | --- |
| Both parents validated (useable as trios) | 9 | 1524 | 1196 | 470 |
| One parent validated (useable as duos) | 19 | 532 | 715 | 684 |
| Missing data for one parent and the other parent was validated (useable as duos) | 38 | 33 | 137 | 122 |
| Neither parent validated | 43 | 197 | 361 | 765 |
| Missing data for one parent and the other parent was not validated | 78 | 33 | 97 | 44 |
| Missing data for both parents | 54 | 0 | 0 | 4 |

Summary of the number of individuals with validated parents across the four breeding groups. An individual's data is labeled "missing" when GBS sequence data was not available for that individual or when replicates could not be resolved with BIGRED.

### Calling and filtering genotypes

Single nucleotide polymorphisms (SNPs) were called from the raw GBS data using pedigree information to select accurate genotypes by calculating genotype posterior probabilities for each individual at each site. The GBS data give the observed counts of each of the two alleles in each individual at each biallelic site: 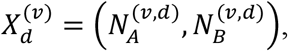 where 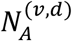 and 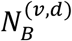 denote the observed counts of allele A and allele B, respectively, in individual *d* at site *v*. Given observed data 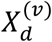 for individual *d* at site *v* and taking the sequencing error rate to be *e* = 0.01, the likelihood for genotype 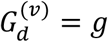 was calculated using the following equation:

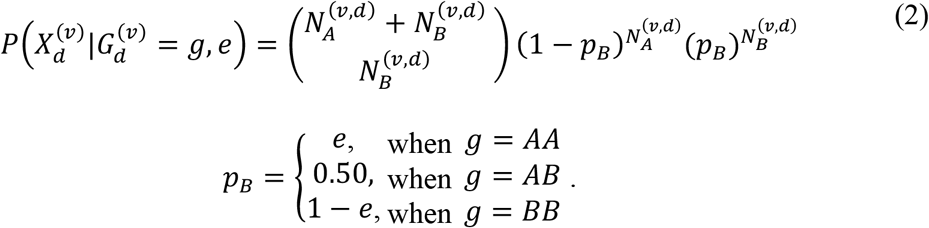

The posterior probabilities for the three genotypes were estimated using the likelihoods defined above with a genotype prior. The genotype prior for each individual *d* was calculated based on the posterior probability genotype distributions of its known parents, following rules of Mendelian inheritance. If individual *d* had only one validated parent or was a founder with no validated parents, its genotype prior for site *v* was calculated using the estimated frequency of the reference allele at site *v* in the population and assuming Hardy-Weinberg Equilibrium (HWE). Genotypes were called for individual *d* at site *v* only if one of the three possible genotypes had a posterior probability greater than or equal to 0.99. We note this method requires calculation of posterior genotype probabilities in a sequential manner, propagating information down the pedigree to subsequent generations.

The dataset was then filtered to remove monomorphic and singleton sites, and sites with more than 30% missing data. The 30% missingness threshold was selected as a compromise between the power to detect crossovers amid noise of poor-quality markers and the resolution affected by the number of markers retained in the dataset. Supplementary Table 1 lists the number of sites retained after applying the filters, ranging from 1,114 to 3,739 SNPs per chromosome, for a total of 35,127 SNPs across the genome.

### Inferring recombination events with SHAPEIT2 and duoHMM

The softwares SHAPEIT2 (Delaneau et al. 2013) and duoHMM were used to phase and impute genotypes, correct switch errors (SEs), and detect intervals surrounding inferred recombination events, following the methods of O’Connell *et al.* 2014. First, phased haplotypes were inferred with SHAPEIT2 without explicit family information, and then the verified pedigrees were used to correct SEs using duoHMM. The duoHMM Hidden Markov Model (HMM) is described in detail in O’Connell *et al.* 2014. Briefly, duoHMM infers the true inheritance states from the observed, imperfect parental and progeny haplotypes. After estimating parameters of the HMM using the Forward Backward algorithm, duoHMM finds the most likely state sequence using the Viterbi algorithm. When duoHMM infers a SE in the Viterbi sequence in either the parent or child, it corrects the haplotypes by switching the phase of all loci following the SE. The algorithm applies these corrections sequentially down through each pedigree.

These steps were carried out internally within SHAPEIT2 by using the ‘—duohmm’ flag. The set of genotypes, verified pedigree information, and a genetic map generated by interpolating genetic distances for the locations of our GBS markers using ICGMC’s composite genetic map were input to produce a haplotype graph encapsulating uncertainty about the underlying haplotypes. SHAPEIT2 was run with 14 burn-in iterations, 16 pruning iterations, and 40 main iterations, with 200 conditioning states per SNP. A window size of 5 Mb was used, based on the developers’ finding that it was advantageous to use a window size larger than 2 Mb when large amounts of identical by descent (IBD) sharing are present (O’Connell *et al.* 2014). The effective population size was set at its default value of 15,000.

After correcting SEs in the SHAPEIT2-inferred haplotypes, duoHMM was run again to infer recombination events. DuoHMM samples a haplotype pair for each individual from SHAPEIT2’s diploid graph and then calculates the probability of a recombination event between markers (O’Connell *et al.* 2014). The inter-SNP recombination probabilities were averaged across ten iterations. A crossover interval was included in subsequent analyses if the interval had an average probability greater than or equal to *t* = 0.5, corresponding to a detection rate of 90.57% and a false discovery rate of 2.89% reported by the developers in simulations with realistic levels of genotyping error (O’Connell *et al.* 2014). Supplementary Figure 1 shows the duoHMM-inferred crossover intervals passing the t=0.5 significance threshold for each chromosome.

### Filtering the SHAPEIT2-duoHMM output

The power to detect recombination events depends on the structure of the pedigree, with most recombination events detectable in a nuclear family with more than two offspring (O’Connell *et al.* 2014). Therefore, those pedigrees consisting of a family with three generations or with more than two offspring were classified as informative towards recombination and were selected for analysis. We refer to the parents of these pedigrees as “informative parents” and the meioses in these pedigrees as “informative meioses”. Of the total 8,678 meioses in the dataset, 7,020 were informative.

### Examining if crossover placements are random and independent events

To examine if crossover placements are random and independent events, the deviance goodness of fit test was used to test whether the distribution of crossovers followed the expected Poisson distribution (Foss et al. 1993; Otto and Payseur 2019). For each chromosome, a Poisson regression was used to model the number of crossovers observed in a given parent-offspring pair as a function of the covariates “parent” and “parental sex,” specifying whether the crossovers were observed in a male or female meiosis. The residual deviance of the regression was used to conduct a chi-square goodness-of-fit test for the model at a Bonferroni-corrected significance level of *α*/*m*, where *α* = 0.05 and *m* = 18 (the number of chromosomes tested).

### Building sex-averaged genetic maps

To build a genetic map for each chromosome, the genetic length of each SNP interval was calculated using the number of recombination events observed in each interval. If a crossover interval spanned multiple SNP intervals, a fraction of the crossover event was assigned to each of the spanned intervals, calculated as 1/(length of the SNP interval). The genetic length of each SNP interval on chromosome *y* was calculated by dividing the number of crossovers in each interval by a scaling factor *n_y_*, where *n_y_* = (the genetic length of chromosome *y* in the ICGMC map)/(the total number of crossovers detected on chromosome *y*), such that the genetic length of each chromosome would be the same as the ICGMC map.

### Examining evidence of sexual dimorphism in crossover rate

To determine whether the distribution of crossover events along each chromosome varied between the sexes, the number of male meiotic crossovers and female meiotic crossovers were compared in 1 Mb windows with a chi-square test in each window. To calculate the expected number of male and female crossovers in a given window, the proportions of informative meioses that were male (0.487) and female (0.514) were multiplied by the total number of crossovers observed in the window. The last window of each chromosome was excluded since it was shorter than 1 Mb. Four of the 510 windows had one or more classes with an expected frequency count of less than five and so were excluded. Each window was tested at a Bonferroni-corrected significance level of *α/N*, where *α* = 0.05 and *N* = 506 (the total number of windows tested). A chi-square test was also conducted genome-wide at a significance level of *α* = 0.05. All chi-square tests were conducted with the *chisq.test()* function in R.

We note that because the dataset contains multiple meioses observed for a single individual, and multiple crossovers counted from each meiosis, the independence assumption of the chi-square test is not perfectly met. However, we expect that this has minimal effect on the interpretation of these tests, due to the size of the population and the magnitude of the observed effect size. In addition, the precedence for the use of chisquare tests for testing heterochiasmy with crossover counts is established in the literature (Drouaud et al. 2007; Kianian et al. 2018; Capilla-Pérez et al. 2021).

### Examining recombination patterns in introgressed regions on chromosome 1 and 4

To examine recombination patterns in the *M. glaziovii* introgression regions on chromosomes 1 and 4, the introgression status of informative parents was first classified based on data described in Wolfe *et al.* 2019. Briefly, a set of Introgression Diagnostic Markers (IDMs) across the cassava genome were identified by comparing a panel of non-admixed *M. glaziovii* individuals with a panel of non-admixed *M. esculenta* individuals.

An IDM was defined as a SNP that is either (1) fixed for different alleles between the *M. glaziovii* and *M. esculenta* reference panels or (2) fixed among *M. esculenta* samples but polymorphic in the *M. glaziovii* panel. Wolfe *et al.* calculated the mean *M. glaziovii* allele dosage at IDMs within 250-kb windows for each individual. To classify the introgression status of each individual, the mean *M. glaziovii* allele dosage across all IDM windows was rounded such that mean dosages falling in the range [0,0.5), [0.5,1.5), and [1.5,2] were rounded to 0, 1 and 2, respectively, to represent generally homozygous non-introgressed, heterozygous, and homozygous introgressed genotypes. There was no introgression data available for one individual (TMS13F1079P0007), so it was excluded from analysis. There were no individuals that were homozygous introgressed on chromosome 4.

To test whether individuals with different introgression statuses have different levels of recombination locally and chromosome-wide, chi-square tests of equal counts were performed with a Bonferroni-corrected significance threshold of *α*/*N*, where *α* = 0.05 and *N* = 4 (the number of regions tested). The expected numbers of crossovers for each introgression class were calculated by multiplying the total number of crossover intervals falling within the introgressed region across all meioses by the proportion of informative meioses contributed by individuals of a given introgression status, which was calculated using the total number of informative meioses counted in the crossover datasets for each chromosome (such that failure to detect the obligatory crossover on a given chromosome did not count towards the difference between observed and expected crossover counts genome-wide). The same chi-square analysis was repeated for the non-introgressed portion of chromosomes 1 and 4 to see if introgression status affected recombination frequency in regions of the chromosome outside of the introgressed region itself.

### Building introgression-specific genetic maps

Introgression status-specific genetic maps were constructed for each of the two chromosomes that contain introgressed segments using the set of crossovers detected in individuals of each introgression class (homozygous non-introgressed, heterozygous introgressed, and homozygous introgressed). The maps were built following the same procedure as for the sex-averaged genetic maps, but the introgression maps were scaled such that their weighted average equaled the sex-averaged map. The genetic length of each SNP interval on a given introgression map was calculated by dividing the number of crossovers detected in parents of a given introgression status in a given interval by *n*_y_, the scaling factor defined above, and then multiplied by *m*, where *m* = (the total number of informative meioses across all three introgression statuses)/(the number of informative meioses contributed by individuals of a given introgression status).

## DATA AVAILABILITY

Raw data and results are publicly available on the Cassavabase FTP: https://www.cassavabase.org/ftp/manuscripts/Chan_et_al_2019/. The README file provides a description of each file. Supplementary Figures are available at FigShare. Code used to perform analysis is available at: https://github.com/serenvillwock/cassava-recombination.

## RESULTS

Using SHAPEIT2 and duoHMM with genotype data at 35,127 SNPs for 5,479 individuals in a validated cassava breeding pedigree, a total of 117,128 crossovers were detected from 7,020 informative meioses. These crossover intervals were used to construct a sex-averaged genetic map with a median resolution of 420,366 bp. To examine the recombination patterns in the regions with known introgressions from *M. glaziovii*, introgression dosage-specific genetic maps were also constructed. To compare these maps to the existing ICGMC map, the genetic positions (cM) of our markers and ICGMC’s markers were plotted against physical position (Mb), shown in Figure 2 for chromosomes 1 and 4 and in Supplementary Figure 2 for all chromosomes.

**Figure 2.**
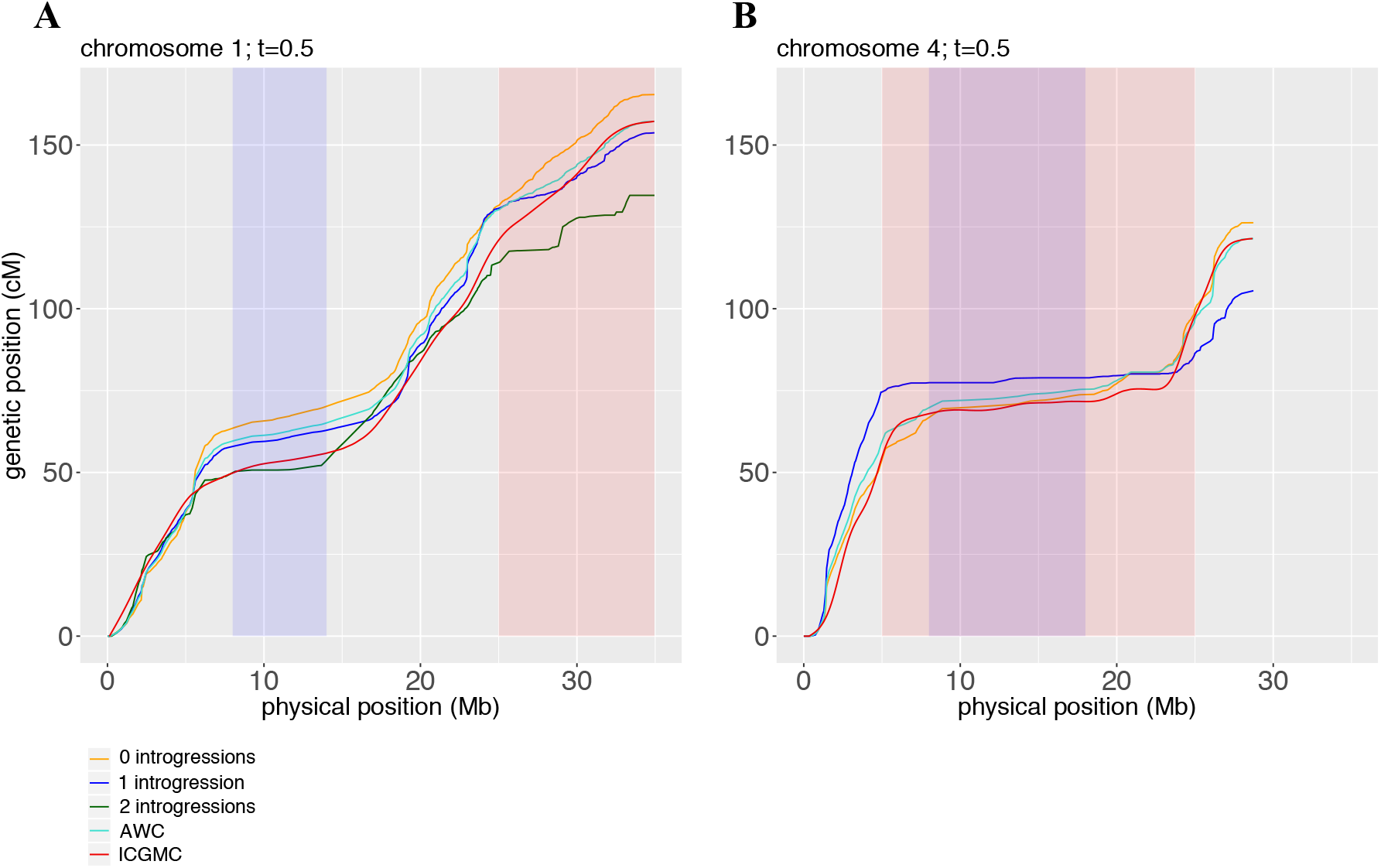
Comparison of constructed genetic maps with the ICGMC map. The genetic position of our GBS markers and ICGMC’s markers are plotted against physical position (Mb) for chromosome 1 (Fig. 2A), and chromosome 4 (Fig. 2B). Centromeric regions are shaded in purple, and *M. glaziovii* introgression regions are shaded in red. Five maps are shown: ICGMC’s map (red), a sex-averaged genetic map constructed from the crossovers detected in all informative parents (turquoise; labeled AWC), and three genetic maps constructed from the crossovers detected in informative meioses of parents that are homozygous non-introgressed (0 introgressions; orange), heterozygous introgressed (1 introgression; blue), and homozygous introgressed (2 introgressions; dark green) in the respective introgression regions on chromosomes 1 and 4. Genetic positions were calculated using the number of crossovers in intervals between SNPs detected by SHAPEIT2-duoHMM passing a significance threshold of t=0.5.

At the qualitative level, the observed crossover distributions were similar to the ICGMC map genome-wide. Both maps showed similar suppression of crossovers in centromeric regions, and the genetic positions generally corresponded well with some regional exceptions, such as in the centromeric region of chromosome 5 (Supplementary Fig. 2). On most chromosomes, there was evidence of crossover interference. In particular, the deviance goodness-of-fit tests were significant at a Bonferroni-corrected significance threshold of 0.00278 for all chromosomes except chromosomes 10, 17, and 18, indicating that crossovers tended to be spaced further apart than would be expected by chance if they were independent events fitting a Poisson model.

A total of 51,357 crossover intervals were identified from 3,446 informative male meioses and 65,771 crossover intervals were identified from 3,574 informative female meioses. The number of crossovers observed genome-wide significantly differed between male and female meioses (chi-square test, p = 5.75 x10^-282^) (Table 3). Females had 10.3% more crossovers than expected if crossover rates were equal between the sexes. The female-to-male ratio of average genome-wide crossovers per meiosis was 1.2.

**Table 3.** Chi-square test results for equal crossover counts between sexes genome-wide.

| Sex | Observed crossover count | Expected crossover count | Chi-square test p-value |
| --- | --- | --- | --- |
| Male | 51357 | 56986 | 5.75 x10<sup>-282</sup> * |
| Female | 65771 | 60142 |  |

Summary of the observed and expected number of crossovers under the null hypotheses of equal recombination rates between the sexes. The observed crossover counts are total genome-wide crossovers detected from meioses of informative parents of each sex. The expected crossover counts were calculated based on the proportion of informative meioses with observed crossovers in parents of each sex.

To investigate variance between the sexes in specific chromosomal regions, chisquare tests for female and male meioses were conducted for crossover counts in 1-Mb windows along each chromosome, shown in Fig. 3 for chromosome 1 and in Supplementary Fig. 3 for all 18 chromosomes. Of the 506 intervals tested, 45 (8.9%) had p-values below the Bonferonni-corrected significance threshold of 9.88 x10^-5^. In these 45 intervals, female crossover count was higher than expected assuming equal male and female crossover rates and male crossover count was lower than expected. Statistically significant intervals were spread throughout the genome and did not consistently appear in any specific region of the chromosomes (Supplementary Fig. 3).

**Figure 3.**
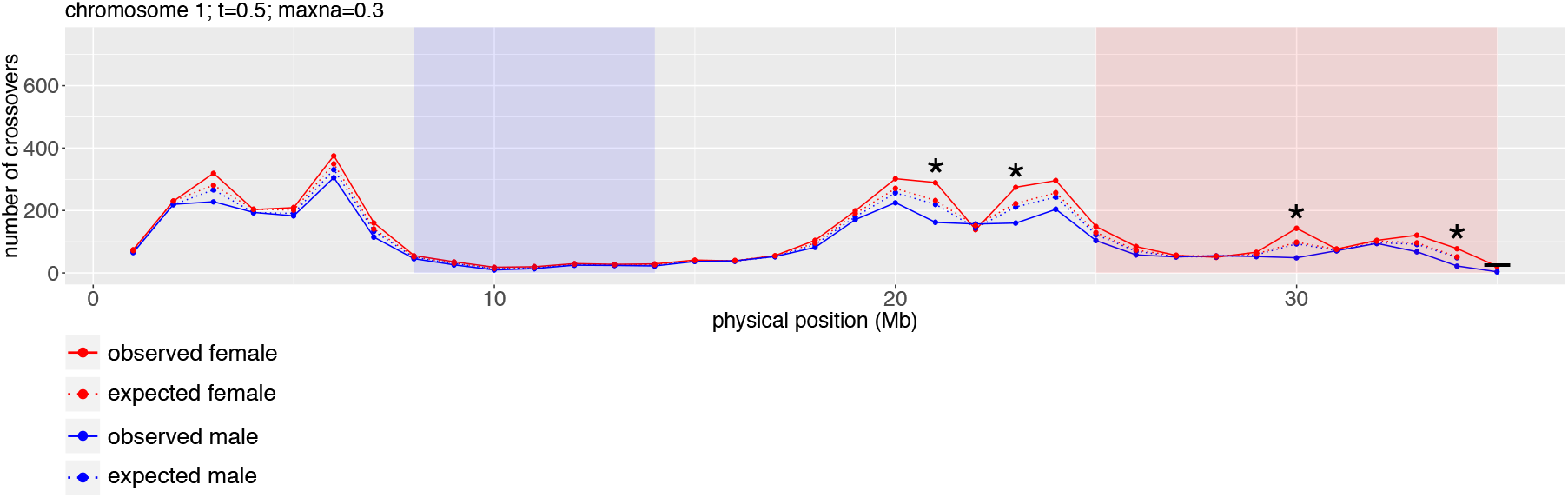
Crossover distribution across chromosome 1 for male and female meioses. The number of crossovers falling within 1 Mb windows are plotted in red for female and blue for male meioses. Solid lines represent observed counts and dashed lines represent expected counts under the null hypothesis of equal recombination frequency in females and males. Asterisks show windows with significantly different crossover frequency between male and female meioses as indicated by a chi-square test at a Bonferonni-corrected α = 0.05. The centromere is highlighted in blue, and the *M. glaziovii* introgression region is highlighted in red. The last window of the chromosome was not tested because it was shorter than 1 Mb (boxed).

To determine whether *M. glaziovii* introgression status affects recombination frequency, chi-square tests were conducted for crossover counts both within and outside of the introgression regions of chromosomes 1 and 4. Table 4 lists the crossover counts that were observed and that were expected under the null hypothesis of equal recombination rates among the introgression classes. At a Bonferroni-corrected significance threshold of 0.0125, the chi-square tests indicated that individuals of different introgression statuses experienced significantly different recombination frequencies within the introgression regions on chromosome 1 (p = 3.97 x10^-14^) and chromosome 4 (p = 6.06 x10^-59^), as well as in the non-introgressed region of chromosome 4 (p = 4.50 x10^-9^), but not in the non-introgressed region of chromosome 1 (p = 3.67 x10^-2^).

**Table 4.** Summary of chi-square test results for equal crossover counts between individuals of different *M. glaziovii* introgression statuses, both within the introgression and in the non-introgressed regions of chromosomes 1 and 4.

| **Chr. 1**<br>**introgression**<br>**status** | **Expected**<br>**count within**<br>**chr. 1**<br>**introgression** | **Observed**<br>**count within**<br>**chr. 1**<br>**introgression** | **Chi-square**<br>**p-value** | **Expected**<br>**count on chr. 1**<br>**outside of**<br>**introgression** | **Observed**<br>**count on chr. 1**<br>**outside of**<br>**introgression** | **Chi-square**<br>**p-value** |
| --- | --- | --- | --- | --- | --- | --- |
| Homozygous<br>non-introgressed | 536 | 680 | 3.97x10<sup>-14</sup> * | 2427 | 2444 | 3.67x10<sup>-2</sup> |
| Heterozygous<br>introgressed | 820 | 698 |  | 3711 | 3744 |  |
| Homozygous<br>introgressed | 89 | 68 |  | 404 | 354 |  |
| **Chr. 4**<br>**introgression**<br>**status** | **Expected**<br>**count within**<br>**chr. 4**<br>**introgression** | **Observed**<br>**count within**<br>**chr. 4**<br>**introgression** | **Chi-square**<br>**p-value** | **Expected**<br>**count on chr. 4**<br>**outside of**<br>**introgression** | **Observed**<br>**count on chr. 4**<br>**outside of**<br>**introgression** | **Chi-square**<br>**p-value** |
| Homozygous<br>non-introgressed | 1212 | 1497 | 6.06x10<sup>-59</sup> * | 2122 | 1986 | 4.50x10<sup>-9</sup> * |
| Heterozygous<br>introgressed | 415 | 130 |  | 726 | 862 |  |

Summary of the number of observed crossovers and those expected under the null hypotheses of equal recombination rates between parental introgression statuses. Observed counts are crossover intervals detected from informative parents of each introgression status that fall within or outside of the respective introgression regions on chromosomes 1 and 4. Expected counts were calculated with the proportions of informative meioses contributed by parents of each introgression status. There were no individuals that were homozygous introgressed on chromosome 4. Asterisks represent significant p values at a Bonferonni-corrected $\alpha = 0.05$ .

In the chromosome 1 introgression region, heterozygous introgressed individuals had 14.5% fewer crossovers than expected under the null hypothesis, while homozygous introgressed individuals had even lower recombination rates (Fig. 2A), with 24.8% fewer crossovers in the introgression region than expected. In the chromosome 4 introgression region, heterozygous introgressed individuals had 68% fewer crossovers than expected, and there were no homozygous introgressed individuals observed. For heterozygous introgressed individuals, more crossovers were observed relative to non-introgressed individuals in the subtelomeric region from 0 to 5 Mb outside of the introgression, but the recombination rate flattened close to zero for most of the chromosome 4 introgression region. In contrast, for non-introgressed individuals, the recombination rate was suppressed only in the centromeric region, but not at either end of the chromosome 4 introgression region that did not overlap the centromeric region (Fig. 2B).

## DISCUSSION

Using IITA’s multi-generational breeding pedigree, a total of 117,128 crossovers were detected and used to construct a new genetic map for cassava, along with a dataset of phased haplotypes, which are improved resources for informing cassava breeding decisions and for studying recombination. In this study, the genetic map was used to investigate (1) sexual dimorphism in crossover number and spatial distribution, and (2) the effect of introgressions from the wild relative *M. glaziovii* on crossover rates.

The genetic map showed that crossover rates vary greatly along the chromosomes. The observations of suppressed recombination in centromeric regions and evidence of crossover interference on most chromosomes were consistent with expectations based on other species (Lawrence et al. 2017). The inferred crossover intervals tended to be longer in the centromeric regions, where there was lower marker density, since a recombination event can only be resolved down to the region between its two flanking heterozygous markers in the parent (Supplementary Fig. 1).

The genetic map shows crossover distributions generally similar to that of the ICGMC map, although it should be noted that information from the ICGMC map was used as input when running SHAPEIT2 and duoHMM. Differences from the ICGMC map could be attributed to several factors. The ICGMC map was generated in 2015 using ten nuclear families with 3480 meioses (ICGMC 2015), while this study used a multi-generational breeding pedigree which had more individuals and more than twice as many informative meioses. In addition, the data used in this analysis were generated using a substantially different variant discovery pipeline and included 35,127 SNPs compared to 22,403 SNPs used by the ICGMC (Chan *et al.,* 2016; ICGMC, 2015). This new genetic map also provides a resource that is directly relevant to cassava breeding programs which use germplasm from the IITA collection.

Certain windows of the genome were identified with significantly more female than male crossovers (Supplementary Fig. 3). This implies that the directionality of crosses matters in cassava, since the female parent of a cross is more likely to recombine than the male parent. Cassava breeders can take advantage of this information to make parent selection decisions, optimizing the chance of finding a new favorable recombination by using an individual as a female parent or the chance of preserving a favorable haplotype by using an individual as a male parent.

The observation of higher crossover rates in female meioses has also been made in other species, including humans (Bhérer et al. 2017) and some other plants (Lenormand and Dutheil 2005), although heterochiasmy in the opposite direction is observed in *A. thaliana* (Drouaud et al. 2007). In maize, the overall number and general distribution of crossovers was found to be similar between the sexes, although higher resolution mapping showed differences in crossover placement relative to specific gene and chromatin features (Kianian et al. 2018). While there were no apparent differences in the overall spatial distribution pattern of crossovers between the sexes in this case, finer differences in crossover position could be investigated with higher resolution mapping.

The mechanism underlying heterochiasmy has been elusive. Lenormand & Dutheil 2005 suggest that heterochiasmy in plants is evolutionarily driven by relative differences in selection pressure on the gametophytes, with less recombination occurring in the sex with greater opportunity for haploid selection. Their observation of lower ratio of male to female recombination rates in plant species with a low selfing rate is consistent with our findings in cassava, an outcrossing species that is rarely selfed in the breeding program. In *A. thaliana* and several animal species, heterochiasmy has been associated with correlated variation in synaptonemal complex length between sexes (Drouaud et al. 2007). Based on observations that transverse filament proteins are necessary for both crossover interference and heterochiasmy in *A. thaliana,* Capilla-Pérez *et al.* 2021 suggest that heterochiasmy is due to interference spacing along synaptonemal complex axes of different lengths in male and female meiocytes.

Interestingly, a low-resolution map of the cassava genome constructed with SSR markers in 2001 showed that the female genetic map was actually shorter than the male map, with a 1.2 ratio of male to female recombination rate (Mba et al. 2001; Lenormand and Dutheil 2005). However, that genetic map was constructed based on a single biparental cross between a cultivar from Nigeria as the female parent and a cultivar from Colombia as the male parent (Fregene et al. 1997), so the observed heterochiasmy in the opposite direction could be attributed to differences in recombination rates between the African and Latin American germplasm. The male and female genetic maps we have constructed with GBS markers across a multigenerational pedigree have higher resolution and are more likely to be representative of the IITA breeding germplasm as a whole. However, disparity in the direction of heterochiasmy has been observed even within the same species, for example between mice subspecies (Dumont and Payseur 2011), so we cannot rule out the possibility that different subpopulations of cassava could vary in which sex exhibits the higher crossover rate.

For both *M. glaziovii* introgression regions on chromosomes 1 and 4, individuals with one or two copies of the introgression showed significantly fewer crossovers within the introgressed regions. These findings are in agreement with previous studies in cassava that have characterized strong LD and lower recombination in the introgression regions relative to the rest of the genome (Rabbi et al. 2017; Wolfe et al. 2019). Evidence of suppressed recombination in introgression regions has also been previously reported in other interspecific hybrids, including in grape (Delame et al. 2019) and tomato (Liharska et al. 1996; Canady et al. 2006). This has important implications for the practicality of introgressing traits from wild germplasm into elite varieties, since linkage drag is exacerbated by low recombination.

A leading hypothesis is that suppressed recombination in the introgression region is due to divergence between the *esculenta* and *glaziovii* haplotypes. Previous studies in plants and animals have associated higher levels of polymorphism between homologs with lower crossover frequency, thought to be due to the anti-crossover role of mismatch repair complexes that recognize interhomolog polymorphism as mismatches during strand invasion (Kolas et al. 2005; Lawrence et al. 2017; Serra et al. 2018). Structural variations in the heterozygous state can especially inhibit crossovers. In *A. thaliana,* crossovers were found to be suppressed within and around the region of inversions and translocations regardless of length (Rowan et al. 2019). In the case of paracentric inversions, crossovers within the inversion can produce acentric and dicentric chromosomes, leading to inviable gametes (McClintock 1931). A paracentric inversion within the chromosome 4 introgression could thus explain why fewer crossovers are observed in that region. While structural polymorphisms in the introgression region have been previously hypothesized, they have yet to be identified (Wolfe et al. 2019).

In addition, the introgression statuses of informative parents were characterized with the average *M. glaziovii* allele dosage in the introgression region rounded to 0, 1, or 2. Individuals classified as generally homozygous introgressed still have some heterozygosity in parts of the introgression region. Therefore, residual interhomologue polymorphisms could be acting to suppress recombination in the introgression even in individuals with an *M. glaziovii* allele dosage of 2.

While interhomologue polymorphism may be involved to some extent in local crossover suppression in the introgression regions, our observations suggest the presence of crossover modifying variants. On chromosome 1, homozygous introgressed individuals have even lower recombination rates that heterozygous introgressed individuals, which implies that a dosage effect of a variant in the introgression rather than solely heterology between homologs is responsible for crossover suppression. On chromosome 4, there were no homozygous introgressed individuals available to determine whether there is a similar dosage effect of the chromosome 4 introgression. However, crossover suppression observed in the heterozygous introgressed state extended even to the non-introgressed region on chromosome 4. Rowan *et al.* 2019 found that crossovers suppression can extend up to 10 kb beyond the border of inversions in *A. thaliana,* but in this case, the observed chromosome-wide crossover suppression is greater than the local or regional suppression that would be expected due to heterozygous polymorphisms alone. Further investigations are needed to test these hypotheses about the mechanism underlying crossover suppression in the introgression regions.

The frequency of the *M. glaziovii* introgression segments in the IITA breeding germplasm has been increasing due to selection on traits that are positively influenced by the introgressions, like root number and dry matter content, although there is also evidence that the introgressions are deleterious in a homozygous state (Wolfe *et al.* 2019). Suppressed recombination in the introgression region limits the ability to purge deleterious load carried along by linkage drag. In the case of the chromosome 4 introgression with very few crossovers in the introgressed region, tightly linked genes may be inherited together as supergenes, which may affect the structure and evolution of cassava populations (Schwander et al. 2014). With the introgression dosage-specific genetic maps, cassava breeders now have a tool to predict the frequency of recombination in the introgression region and plan population sizes accordingly to increase the chance of finding a desired recombination.

## ACKNOWLEDGEMENTS

We acknowledge the UK’s Foreign, Commonwealth & Development Office (FCDO) and the Bill & Melinda Gates Foundation (Grant INV-007637 http://www.gatesfoundation.org) for funding this work through the “Next Generation Cassava Breeding Project”. We thank the entire Next Generation Cassava Breeding team for contributing to this study in the field and lab. We give special thanks to the International Institute of Tropical Agriculture (IITA), Ibadan, Nigeria for sharing their pedigree records with us for this study. Marnin Wolfe shared datasets on *M. glaziovii* introgressions with us, for which we are grateful. A.L.W. was supported by NIH grant R35 GM133805.

